# Strain-specific differences in the interactions of the cucumber mosaic virus 2b protein with the viral 1a and host Argonaute 1 proteins

**DOI:** 10.1101/2023.08.31.555694

**Authors:** Sam Crawshaw, Lewis G. Watt, Alex M. Murphy, John P. Carr

**Affiliations:** Department of Plant Sciences, University of Cambridge, Downing Street, Cambridge CB2 3EA

**Keywords:** small RNA-binding, RNA silencing, viral host manipulation, symptom induction, aphid vector, subcellular localization

## Abstract

The cucumber mosaic virus (CMV) 2b protein is a potent counter-defense protein and symptom determinant that inhibits antiviral silencing by titration of short double-stranded RNAs. Expression of the 2b protein from the CMV Subgroup IA strain Fny-CMV in transgenic *Arabidopsis thaliana* plants disrupts microRNA-mediated cleavage of host mRNAs by binding ARGONAUTE 1 (AGO1), leading to symptom-like phenotypes. This also triggers AGO2-mediated resistance against CMV and strong resistance to CMV’s aphid vectors, which would be deleterious to viral fitness. However, in authentic viral infections the Fny-CMV 1a protein modulates 2b-AGO1 interactions, which inhibits induction of AGO2-mediated virus resistance and resistance to aphid vectors. Contrastingly, the 2b proteins encoded by the Subgroup II LS-CMV strain or the recently discovered Subgroup IA strain Ho-CMV induce no apparent symptoms. Confocal laser scanning microscopy, bimolecular fluorescence complementation and co-immunoprecipitation showed that the Fny-CMV and Ho-CMV 2b proteins interact with the Fny-CMV and LS-CMV 1a proteins whilst the CMV-LS 2b protein does not. However, the Fny-CMV, Ho-CMV and LS-CMV 2b proteins all interacted with AGO1, but while AGO1-Fny2b complexes occurred in the host cell nucleus and cytoplasm, the corresponding AGO1-2b complexes for LS-CMV and Ho-CMV accumulated almost exclusively in nuclei. *AGO2* transcript accumulation was used to assess the inhibition of AGO1-mediated miRNA-regulated mRNA cleavage. While Fny-CMV 2b induced a five-fold increase in *AGO2* accumulation, the LS-CMV and Ho-CMV 2b proteins induced only two-fold increases. Thus, these 2b proteins bind AGO1 but are less effective at inhibiting AGO1 activity. We conclude that the intracellular localization sites of 2b-AGO1 complexes influences the degree to which a 2b protein can inhibit microRNA-mediated host mRNA degradation and that cytoplasmic AGO1 has the strongest influence on miRNA-mediated cellular mRNA turnover.

## INTRODUCTION

*Cucumber mosaic virus* (CMV), the type species of the genus *Cucumovirus* in the family *Bromoviridae*, is an agronomically important virus that is vectored by over 80 species of aphid and infects an estimated 1,071 species of plants across 521 genera (Yoon et al., 2019). CMV has a tripartite positive-sense RNA genome that encodes five viral proteins: the 1a protein, which has methyltransferase and RNA helicase active sites, and is translated from genomic RNA1; the 2a RNA-dependent RNA polymerase protein, which is translated directly from genomic RNA2; the multifunctional 2b protein, which is best known as a viral suppressor of RNA silencing (VSR), which is translated from a sub-genomic RNA (RNA4A) derived from RNA2, and the movement protein and coat protein that are, respectively, translated from genomic RNA 3 and the RNA3-derived sub-genomic RNA4 (Palukaitis and García-Arenal, 2003; Li and Ding, 2006; Jacquemond, 2012). CMV strains are classified into the Subgroups IA, IB and II based on RNA sequence similarity (Palukaitis and García-Arenal, 2003; Balaji et al., 2008; Jacquemond, 2012).

In *Arabidopsis thaliana*, a natural host for CMV (Pagán et al., 2010), Subgroup II CMV strains generally cause mild or unnoticeable symptoms, compared to the symptoms induced by strains of Subgroups IA and IB (Mochizuki and Ohki, 2011). For example, the Subgroup IA Fny-CMV strain causes more severe symptoms in *A. thaliana* accession Col-0 than the Subgroup II strains LS-CMV and Q-CMV (Lewsey et al., 2007). However, symptom severity is not always dictated by Subgroup, as demonstrated by the recently characterized Subgroup IA strain Ho-CMV isolated from the perennial wild plant *A. halleri* (Takahashi et al., 2022). Ho-CMV, despite accumulating to similarly high levels in *A. thaliana* as Fny-CMV, causes no symptoms. Ho-CMV shows high RNA sequence similarity to Fny-CMV, for example, their 1a open reading frames share 97% amino acid sequence identity. However, the most striking contrast between the sequences of Ho-CMV and Fny-CMV are four differences in the amino acid sequences of their respective 2b proteins.

The CMV 2b protein is an important determinant of symptom severity in several plants including *A. thaliana* (Ding et al. 1996; Shi et al. 2002, 2003; Du et al. 2007; Lewsey et al., 2007, 2009; Ziebell et al. 2007; Mochizuki and Ohki, 2011). In transgenic *A. thaliana*, constitutive expression of the 2b protein of Fny-CMV (Subgroup IA) induces strong symptom-like phenotypes while expression of 2b proteins from Subgroup II strains LS-CMV and Q-CMV does not (Lewsey et al., 2007). Surprisingly, constitutive expression of the 2b protein of the Subgroup IA Ho-CMV did not modify the phenotype of transformed *A. thaliana* when compared to control transformed plants (Takahashi et al., 2022). Thus, the differences in 2b amino acid sequence of the two Subgroup IA viruses, Ho- and Fny-CMV, may be important in the contrasting severity of symptoms seen in *A. thaliana*.

In CMV-infected plants and in *2b*-transgenic plants, the respective severity of symptoms or of phenotype are linked to the ability of different CMV strains’ 2b proteins to disrupt the regulation of host gene expression by microRNAs (miRNAs) (Zhang et al., 2006; Lewsey et al., 2007; Siddiqui et al., 2008; Du et al., 2014b). This is thought to be in large part due to inhibition of the activity of the host RNA silencing factor, Argonaute 1 (AGO1). AGO1 is an endonuclease that uses miRNAs and short-interfering RNAs as guides to direct site-specific cleavage of RNA molecules at complementary sequences or to inhibit mRNA translation on ribosomes (Baumberger and Baulcombe, 2005). Recent work showed that the 2b-AGO1 interaction is modulated by a competing interaction of the 2b protein with the CMV 1a protein, combined with re-localization of the 1a-2b protein complexes to P-bodies (Watt et al., 2020). It was suggested that in *A. thaliana*, the severe symptoms induced by Subgroup IA strains such as Fny-CMV can be explained by the interaction of the 2b protein with AGO1, whereas the mild symptoms exhibited by Subgroup II strains such as LS-CMV and Q-CMV result from a lack of any interaction between the 2b proteins of these strains (which are identical in amino acid sequence) with AGO1 (Lewsey et al. 2007). Intriguingly, it was reported that the 2b protein of the Subgroup IA Ho-CMV strain does not interact with AGO1 and this may explain the lack of symptoms in *A. thaliana* plants (Takahashi et al., 2022). In this study we investigated the 2b proteins of the CMV strains Fny, Ho and LS with respect to their subcellular localization and contrasting abilities to interact with AGO1 and with the CMV 1a protein.

## RESULTS

### Partitioning of the Ho-CMV 2b between the cytoplasmic and nucleus is intermediate between that of the 2b proteins of Fny-CMV and LS-CMV

The Ho-CMV 2b protein amino acid sequence differs from its Fny-CMV orthologue at four residues (Figure 1). The presence of alanine at residue 72 of the Ho-CMV 2b protein differs from the valine present at this position in the Fny-CMV 2b protein and the majority of orthologues encoded by Subgroup IA and IB strains although this renders it similar to the 2b protein of another Subgroup IA virus, Y-CMV, at this position. The proline at residue 80, instead of a serine as in most other Subgroup IA and IB CMV strains, also marks a similarity between the Ho-CMV and Y-CMV 2b orthologues (Figure 1). Potentially the most significant residues that distinguish the Ho-CMV 2b protein with respect to symptom expression, are the alanine residue at position 47, which replaces a highly conserved serine, and the valine residue at position 106, which replaces a highly conserved alanine residue, both of which are conserved across Subgroup IA, IB and II CMV strains.

**Figure 1.**
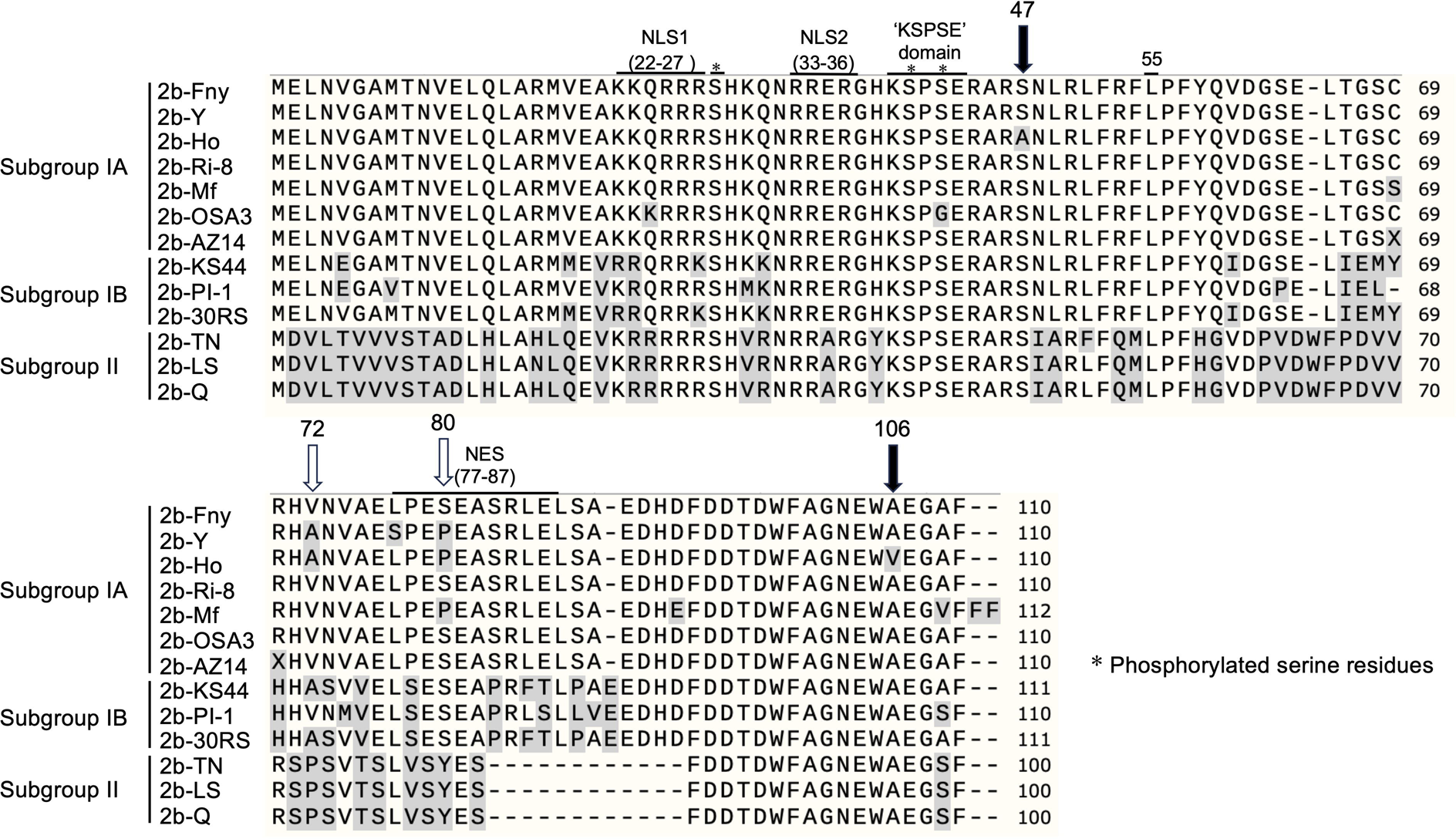
The amino acid sequences of 2b proteins encoded by a selection of strains of CMV from Subgroups IA, IB and II. The Fny-CMV 2b protein sequence is given as a reference to compare with 2b protein sequences from a small number of other CMV strains to highlight certain differences and similarities between 2b proteins of viruses in Subgroups IA, IB and II. The nuclear localisation sites (NLS1 and NLS2), nuclear export site (NES), self-interaction site (residue 55) are over-lined, known phosphorylation sites (including the ‘KSPSE domain’) are indicated with asterisks, while differences between the Ho-CMV 2b protein sequence and that of the Fny-CMV 2b protein are indicated by white arrows at residues where there is variation between different strains (residues 72 and 80) and with solid arrows at residues where there is normally strong inter-strain conservation (residues 47 and 106). Differences in 2b protein primary sequences between the Fny-CMV and other strains are highlighted in grey. The numbering of amino acid residues is based on the Fny-CMV 2b protein sequence. The GenBank accession numbers for the sequences used in this alignment are NC002035 for Fny-CMV (2b-Fny), D12538 for Y-CMV (2b-Y), LC593245 for Ho-CMV (2b-Ho), AM183118 for RI-8-CMV (2b-RI-8), AJ276480 for Mf-CMV (2b-Mf), HE971489 for OSA3-CMV (2b-OSA3), QBH72281 for AZ14-CMV (2b-AZ14), CBG76802 for KS44-CMV (2b-KS44), CAJ65577 for PI-1-CMV (2b-PI-1), FN552601 for 30RS-CMV (2b-30RS), BAD15371 for TN-CMV (2b-TN), AF416900 for LS-CMV (2b-LS) and Q66125 for Q-CMV (2b-Q).

Cucumoviral 2b proteins contain sequences that control their localization in host cell nuclei and their ability to dimerize (Xu et al., 2013; Gonzalez et al., 2010) (Figure 1). To examine the localisation pattern of 2b proteins, sequences encoding RFP tags fused to the C-termini of the 2b protein were inserted into T-DNA constructs and transiently expressed in leaves of *Nicotiana benthamiana* plants. Consistent with previous localization and fractionation studies, the LS-CMV 2b protein was highly concentrated in the nucleus, while the Fny-CMV 2b protein was abundant not only in the nucleus but also across the cytoplasm and cytoskeleton (Figure 2A) (Mayers et al., 2000; Gonzalez et al., 2010). Ho-CMV 2b protein accumulation was not restricted to the nucleus but a greater proportion accumulated there compared to the distribution of the Fny-CMV 2b protein (Figure 2A).

**Figure 2.**
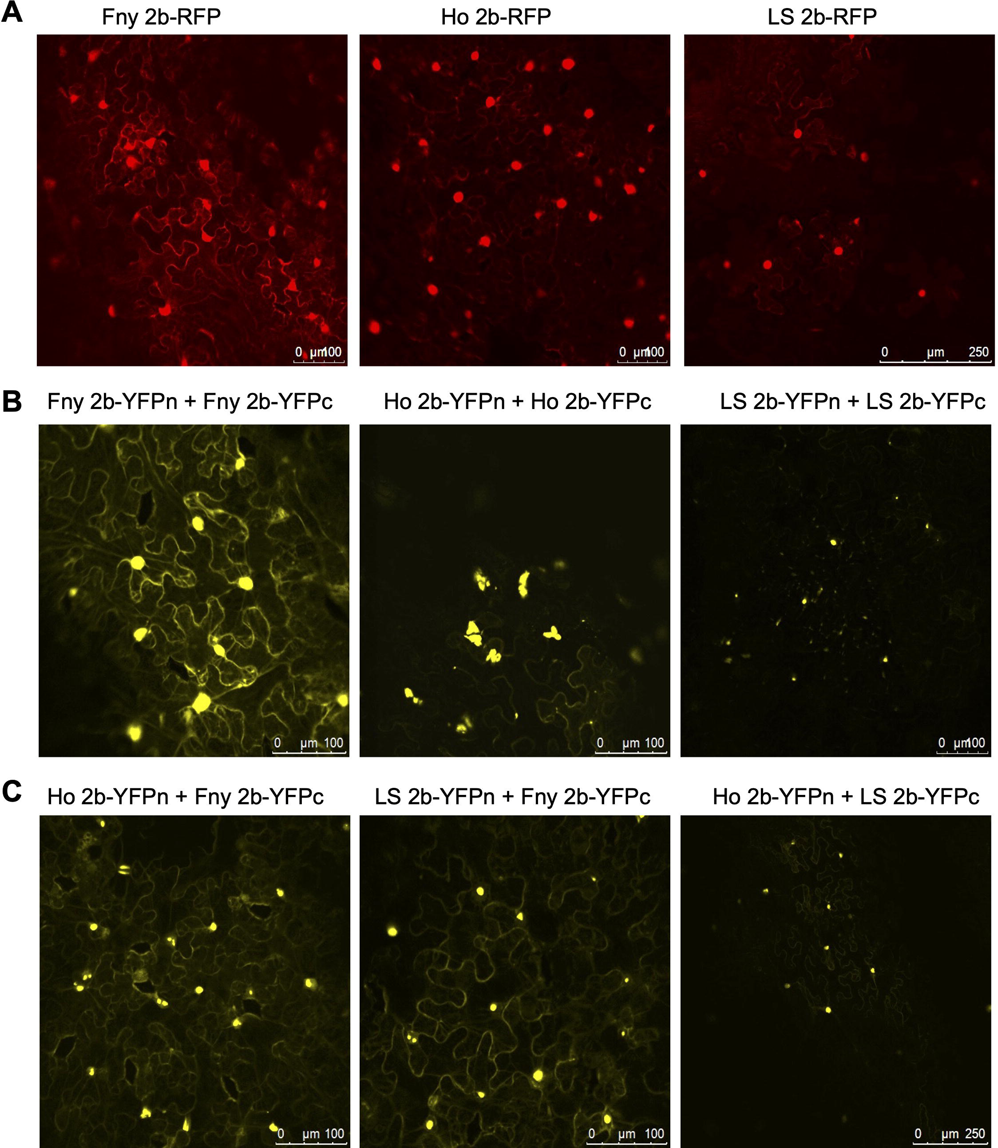
Subcellular localization of cucumber mosaic virus 2b proteins from different CMV strains. (A) Agroinfiltration was used for transient expression in *N. benthamiana* leaves of C-terminal red fluorescent protein (RFP)-2b fusion protein derived from the 2b proteins of Fny-CMV (Fny 2b-RFP), LS-CMV (LS 2b-RFP), or Ho-CMV (Ho 2b-RFP). **(B)** Bimolecular fluorescence complementation was used to compare the self-interaction properties of the 2b proteins of Fny-CMV, LS-CMV, and Ho-CMV, using fusion proteins with the N- and C-terminal domains of the yellow fluorescent protein (2b-YFPn and 2b-YFPc). The 2b proteins of all three strains formed homodimers in vivo, indicated by yellow fluorescence. The intracellular distributions of these homodimers were consistent with those seen in panel A for 2b-RFP proteins and show a greater proportion of the Fny-2b protein being present in the cytoplasm. (**C**) Bimolecular fluorescence complementation showed that the 2b proteins of Fny-CMV, LS-CMV, and Ho-CMV were capable of heterodimerization. When 2b-YFPc proteins derived from LS-CMV or Ho-CMV were co-expressed with 2b-YFPn derived from Fny-CMV strain, fluorescence was visible in the cytoplasm as well as the nucleus. This suggests that the localization behavior of the Fny-CMV 2b protein is dominant over that for LS-CMV 2b and Ho-CMV 2b proteins.

To examine the localization pattern of self-interacting 2b proteins, sequences encoding N-and C-terminal split yellow fluorescent protein (YFP) tags fused to the C-termini of the 2b protein were inserted into T-DNA constructs and transiently expressed in leaves of *N. benthamiana* plants. Bimolecular fluorescence complementation (BiFC) was used to detect the formation of 2b protein homodimers (Figure 2B). We observed that the Fny-CMV 2b protein formed dimers which is consistent with previous work (Chen et al., 2008; Gonzalez et al., 2010; Xu et al., 2013). We also showed that 2b proteins from Ho-CMV and LS-CMV were able to homodimerize (Figure 2B). To our knowledge, this is the first time that dimerization has been shown for a subgroup II 2b protein. The formation of homodimers in the 2b proteins of all three strains was expected since the leucine residue at position 55, which is thought to control dimerization (Xu et al., 2013), is present in all three orthologs (Figure 1). The localization pattern of self-interacting sYFP-tagged 2b proteins was the same as observed for RFP-tagged 2b proteins with Fny-CMV 2b proteins having a greater cytoplasmic presence than 2b proteins from LS-CMV or Ho-CMV. We also showed that the Fny-CMV 2b protein was able to form heterodimers with the LS-CMV and Ho-CMV orthologs (Figure 2C). The subcellular distribution of these heterodimers was similar to that observed for homodimers of the Fny-CMV 2b protein (Figure 2B). This suggests that the Fny-CMV 2b protein is dominant over the 2b proteins of the two other strains in determining the subcellular accumulation pattern.

### The interaction of CMV 2b and 1a proteins is Subgroup-specific

The Fny-CMV 1a and 2b proteins physically interact (Watt et al., 2020). We investigated if the Fny-CMV 1a protein can interact with 2b proteins of other CMV strains. First, we imaged the subcellular localization of the Fny-CMV 1a protein, C-terminally fused with either GFP or RFP sequences, and GFP-fused or RFP-fused 2b proteins, using agroinfiltration to introduce T-DNA constructs encoding these proteins into patches of *N. benthamiana* leaf tissue (Figure 3). Consistent with previous work, the localization of the Fny-CMV 2b protein in the nucleus and cytoplasm (Figure 2) was altered so that it accumulated together with a proportion of the 1a protein in specks (Figure 3) that were previously shown to relate to P-bodies (Watt et al., 2020). In contrast, the predominantly nuclear localization of LS-CMV 2b protein (Figure 2) was unaffected by co-expression with the Fny-CMV 1a protein (Figure 3). Only a small proportion of Ho-CMV 2b protein co-localized with the Fny-CMV 1a protein into P-bodies (Figure 3).

**Figure 3.**
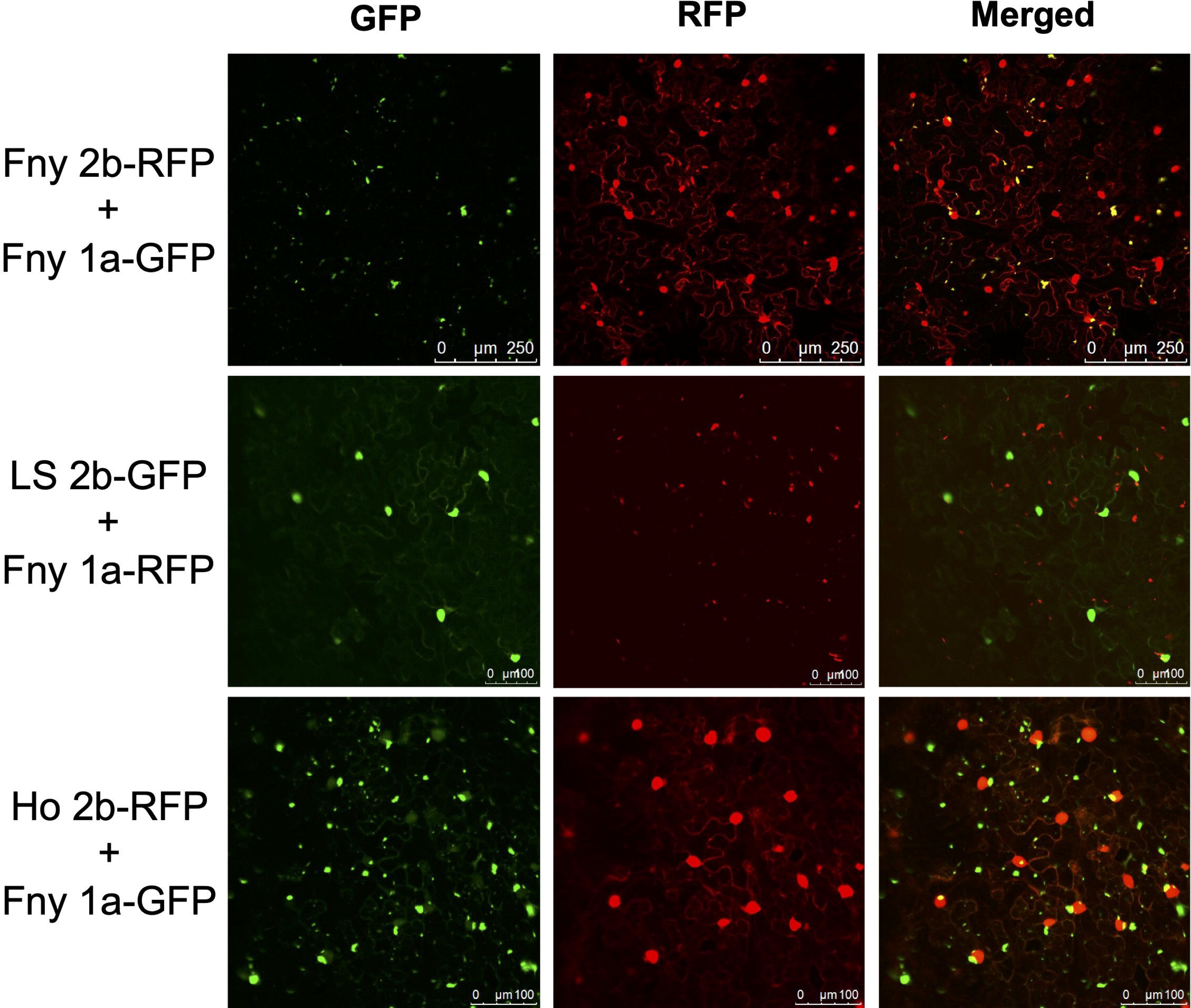
Subcellular localization of the 2b proteins of three cucumber mosaic virus strains and the Fny-CMV 1a protein. Using agroinfiltration, C-terminal RFP or GFP fusion proteins derived from the 2b proteins of Fny-CMV (Fny 2b-RFP), LS-CMV (LS 2b-GFP), or Ho-CMV (Ho 2b-RFP), and the Fny-CMV 1a protein (Fny 1a-GFP or Fny 1a-RFP) were co-expressed in *N. benthamiana* leaves in the combinations shown. Fluorescent signals were imaged using confocal scanning laser microscopy. Fny 2b-RFP accumulated in the nucleus and cytoplasm, with a proportion co-localizing with the 1a-GFP (merged signal shown as yellow), consistent with previous results (Watt et al., 2020). The LS 2b-GFP protein accumulated in the nucleus with no detectable co-localization with 1a-RFP. Ho 2b-RFP protein accumulated in the nucleus, with a small portion weakly co-localizing with the 1a-GFP signal in cytoplasmic foci (merged signal shown as yellow).

Co-localization experiments suggested that a direct interaction between the Fny-CMV 1a protein and the LS-CMV 2b protein was unlikely but did not rule out a heterologous interaction between the Fny-CMV 1a protein and the Ho-CMV 2b protein (Figure 3). The possibility of intermolecular interactions between the Fny-CMV 1a protein and the 2b proteins of LS-CMV and Ho-CMV were investigated directly using BiFC (Figure 4) and co-immunoprecipitation assays (Figure 5). For BiFC assays, T-DNA constructs encoding fusion proteins of the N- or C-terminal portions of YFP (sYFPn and sYFPc, respectively) with the respective 2b proteins and the Fny-CMV 1a protein were transiently expressed in *N. benthamiana* (Figure 4). This confirmed that the Fny-CMV 1a and 2b proteins interact *in planta* but no interaction between the 2b protein of LS-CMV with the Fny-CMV 1a protein was detectable (Figure 4A). However, BiFC assays indicated that the 2b protein of Ho-CMV can interact with Fny-CMV 1a protein (Figure 4A). Interestingly, BiFC analyses showed that while the 2b protein of Fny-CMV can localize with and directly interact with both the Fny- and LS-CMV 1a proteins, the LS-CMV 2b protein does not interact with either of these CMV 1a proteins (Figure 4B).

**Figure 4.**
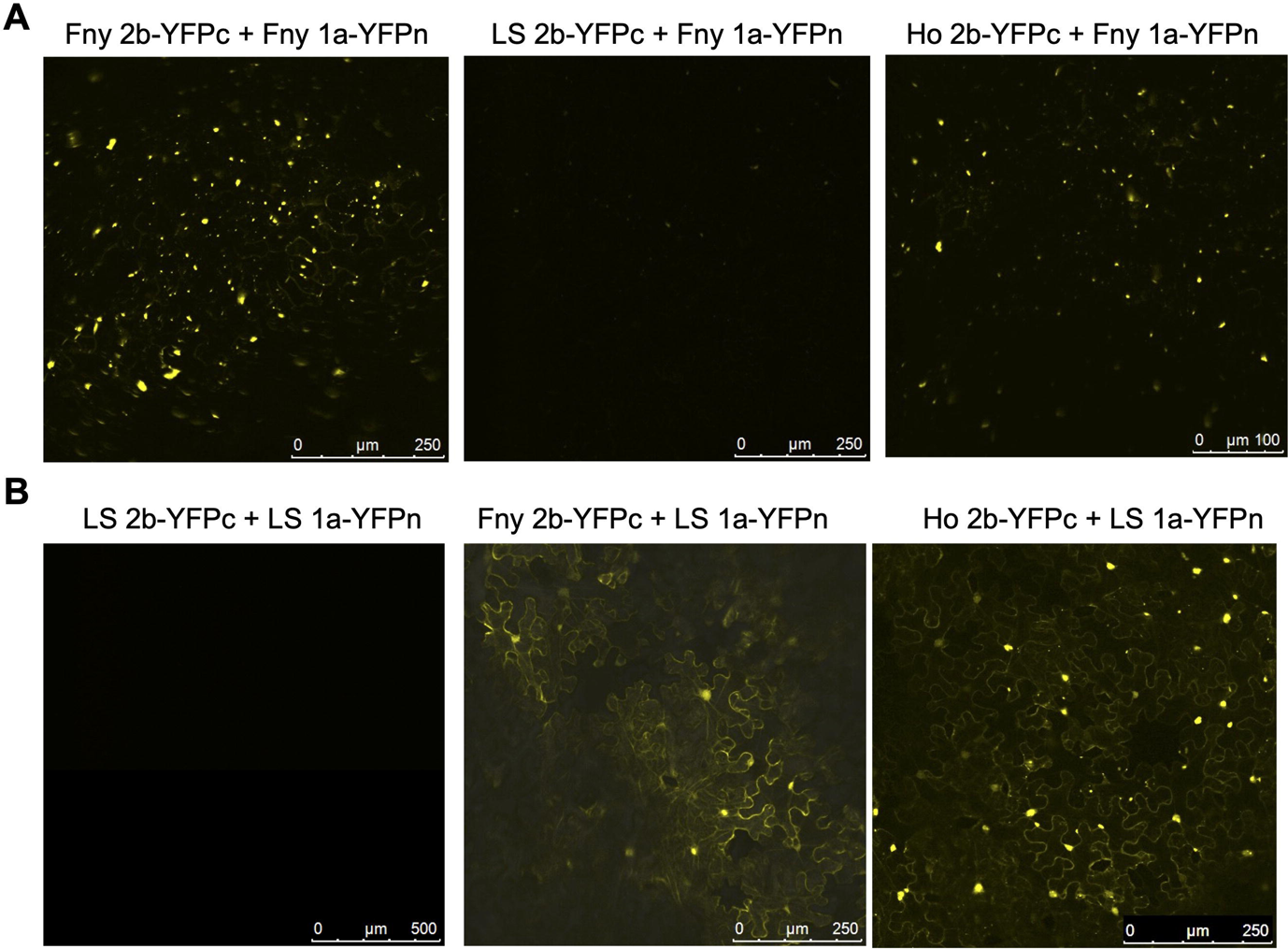
Interactions of the Fny-CMV, LS-CMV and Ho-CMV 2b proteins with the 1a proteins of Fny-CMV and LS-CMV. The 2b protein orthologs of Fny-CMV, LS-CMV and Ho-CMV were fused at their C-termini with the C-terminal domain of split yellow fluorescent protein (respectively: Fny 2b-YFPc; LS 2b-YFPc, and Ho 2b-YFPc). Using agroinfiltration in *N. benthamiana* leaves, these fusion proteins were co-expressed with YFP N-proximal domain fusion proteins with the 1a proteins of Fny-CMV (Fny 1a-YFPn) or LS-CMV (LS 1a-YFPn). Direct protein-protein interactions *in vivo* were revealed by bimolecular fluorescence complementation and resulting fluorescence imaged by confocal laser scanning microscopy. The data show that LS-CMV 2b protein does not physically interact with either the LS-CMV or Fny-CMV 1a proteins, whereas the 2b proteins of Fny-CMV and Ho-CMV can interact with both 1a orthologs.

**Figure 5.**
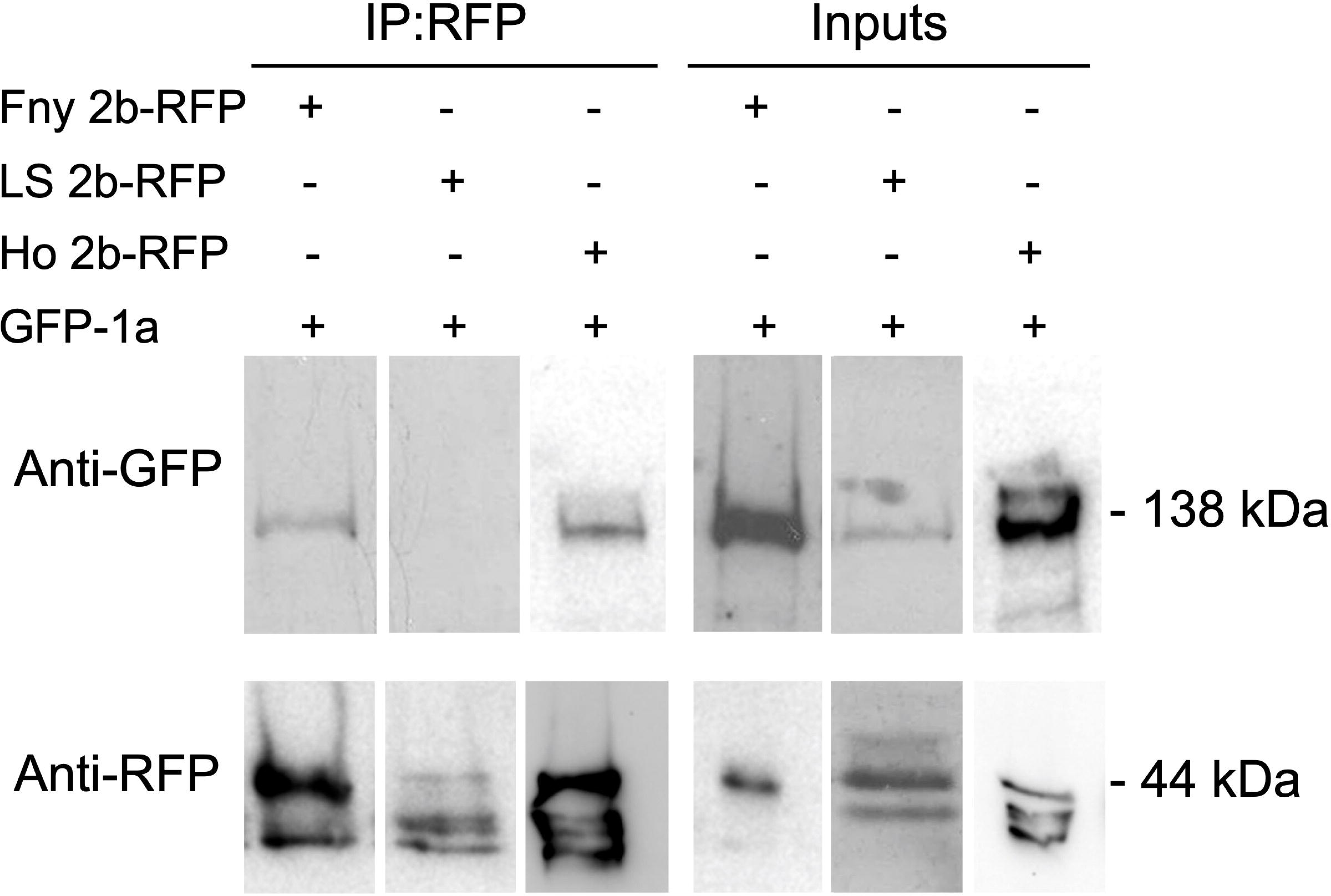
Interactions of the cucumber mosaic virus 1a protein with 2b protein orthologs *in planta* examined by co-immunoprecipitation. Proteins were transiently expressed in *N. benthamiana* leaf tissue using agroinfiltration. *A. tumefaciens* cells carried T-DNAs designed to express a fusion protein of green fluorescent protein and the CMV 1a protein (GFP-1a) mixed with cells carrying T-DNA vectors encoding 2b-RFP fusion proteins engineered from the respective 2b orthologs of Fny-CMV (Fny 2b-RFP), LS-CMV (LS 2b-RFP) and Ho-CMV (Ho 2b-RFP). Total protein was extracted from leaf samples and subjected to immunoprecipitation with RFP-Trap beads (IP:RFP) followed by immunoblot analysis with anti-GFP antibodies to detect GFP-1a fusion proteins. GFP-1a was detected in all input samples with a corresponding band of approximately 138kDa. However, following RFP-pull down, GFP-1a could only be detected when co-expressed with the RFP tagged version of 2b from the Fny or Ho strain of CMV and not the LS strain version. Original blots used to make composite image are shown in Figure S3.

Co-immunoprecipitation assays confirmed that physical interactions occur between the Fny-CMV 1a protein and the 2b protein of Fny-CMV or Ho-CMV but not that of LS-CMV (Figure 5). Transiently expressed GFP-tagged Fny-CMV 1a protein in the presence or absence of 2b-RFP fusion proteins derived from Fny-CMV, LS-CMV or Ho-CMV was immunoprecipitated from leaf homogenates using anti-RFP agarose magnetic beads and analyzed by western blotting using anti-GFP antibodies to detect any 2b proteins complexed with the Fny-CMV 1a protein. The Fny-CMV 1a protein-GFP fusion co-immunoprecipitated with the RFP-tagged 2b protein from Fny-CMV or Ho-CMV but not with the RFP-tagged LS-CMV 2b protein (Figure 5), in line with results from the BiFC assays.

### CMV strain-specific differences in 2b-AGO1 interactions

The Fny-CMV 2b protein is known to interact with AGO1 (Zhang et al., 2006; González et al., 2010), but it is less clear if 2b proteins from Subgroup II strains, such as the identical proteins encoded by LS-CMV and Q-CMV, also interact with this factor. Zhang and colleagues (2006) commented on data (not shown) indicating an interaction between the Q-CMV 2b protein and AGO1 but, to our knowledge, the results were not published. In order to definitively demonstrate that 2b proteins from Subgroup II strains can interact with AGO1, constructs encoding RFP or GFP tagged 2b protein from Fny-CMV, LS-CMV or Ho-CMV were transiently expressed with the corresponding GFP- or RFP-tagged variants of the *A. thaliana* AGO1 protein in *N. benthamiana* leaf tissue. Confocal laser scanning microscopy revealed that the AGO1 protein co-localized with the Fny-CMV, Ho-CMV and LS-CMV 2b proteins (Figure 6). BiFC was used to assess interactions between AGO1 and the 2b proteins and it was apparent that the Fny-CMV, Ho-CMV and LS-CMV 2b proteins all interacted with AGO1 (Figure 7A). This interaction appeared to occur in the nucleus for all three 2b proteins, but for the Fny-CMV 2b protein, the interaction with AGO1 in other parts of the cell was more apparent than for the 2b proteins of LS-CMV and Ho-CMV (Figure 7A, B). Co-immunoprecipitation confirmed that AGO1 interacts with 2b proteins of Fny-CMV, Ho-CMV, and LS-CMV (Figure 8).

**Figure 6.**
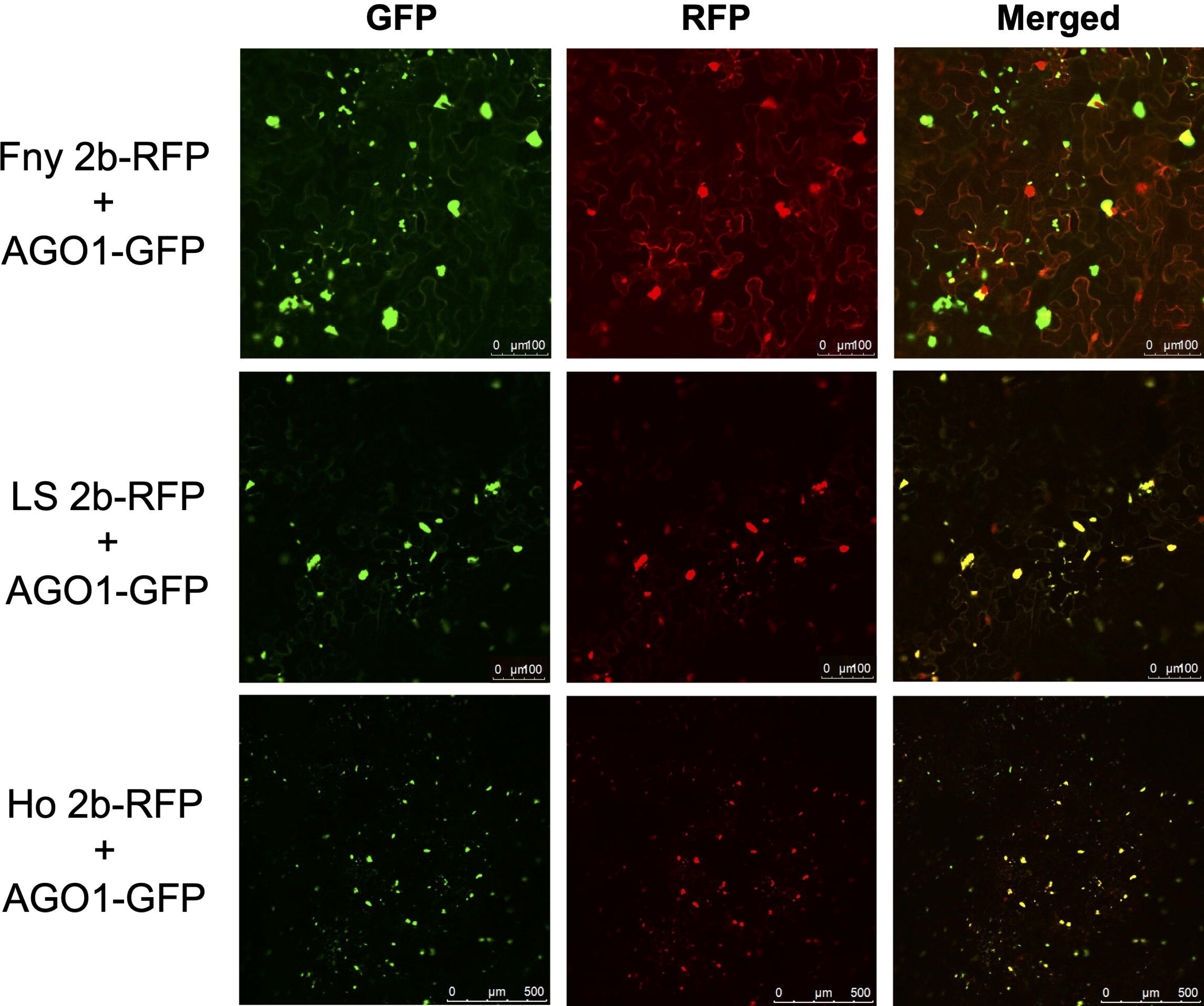
Subcellular localization patterns of Argonaute 1 and the 2b proteins of three cucumber mosaic virus strains. C-terminal RFP fusions of the Fny-CMV (Fny 2b-RFP), LS-CMV (LS 2b-RFP) and Ho-CMV 2b (Ho 2b-RFP) proteins were co-expressed with an Argonaute 1-GFP fusion protein (AGO1-GFP) in *N. benthamiana* leaves using agroinfiltration. Fluorescence was imaged by confocal scanning laser microscopy: a yellow signal in the merged channel indicates co-localization of RFP and GFP signals. A portion of the Fny 2b-RFP signal co-localized with the AGO1-GFP signal in the nucleus, cytoplasm and at cytoplasmic foci. LS 2b-RFP accumulated in the nucleus where it co-localized with nuclear-localized AGO1-GFP, while Ho 2b-RFP accumulated not only in the nucleus but also throughout the cytoplasm.

**Figure 7.**
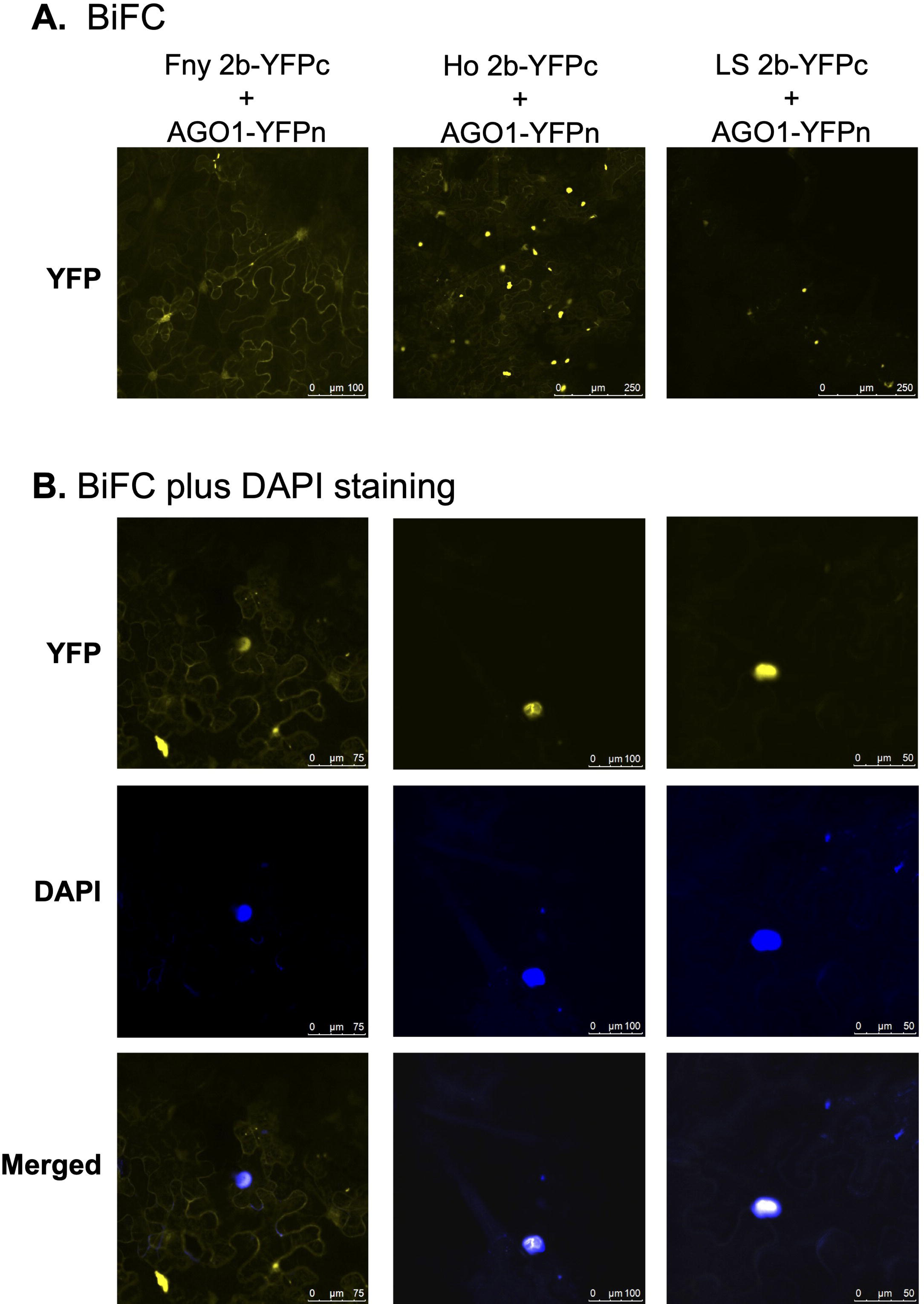
Argonaute 1 interacts not only with the 2b protein of Fny-CMV but also with the 2b orthologs of LS-CMV and Ho-CMV. The 2b proteins encoded by the Fny, LS and Ho strains of CMV were fused at their C-termini with the C-terminal domain of split yellow fluorescent protein (respectively indicated by: Fny 2b-YFPc; LS 2b-YFPc, and Ho 2b-YFPc). Using agroinfiltration these fusion proteins were co-expressed with the Argonaute 1 protein-yellow fluorescent protein N-proximal domain fusion protein (AGO1-YFPn) in *N. benthamiana* leaves. Direct protein-protein interactions *in vivo* were revealed by bimolecular fluorescence complementation and imaged by confocal laser scanning microscopy. (A) Fluorescence indicated that direct interactions occurred between AGO1-YFPn and 2b-YFPc fusion proteins derived from the 2b orthologs of Fny-CMV, LS-CMV and Ho-CMV. (B) The DNA marker DAPI was used to highlight cell nuclei. Imaging confirmed that 2b-AGO1 complexes strongly localized to nuclei but in the case of the Fny-CMV 2b protein there was also easily observable accumulation of 2b-AGO1 complexes in the cytoplasm.

**Figure 8.**
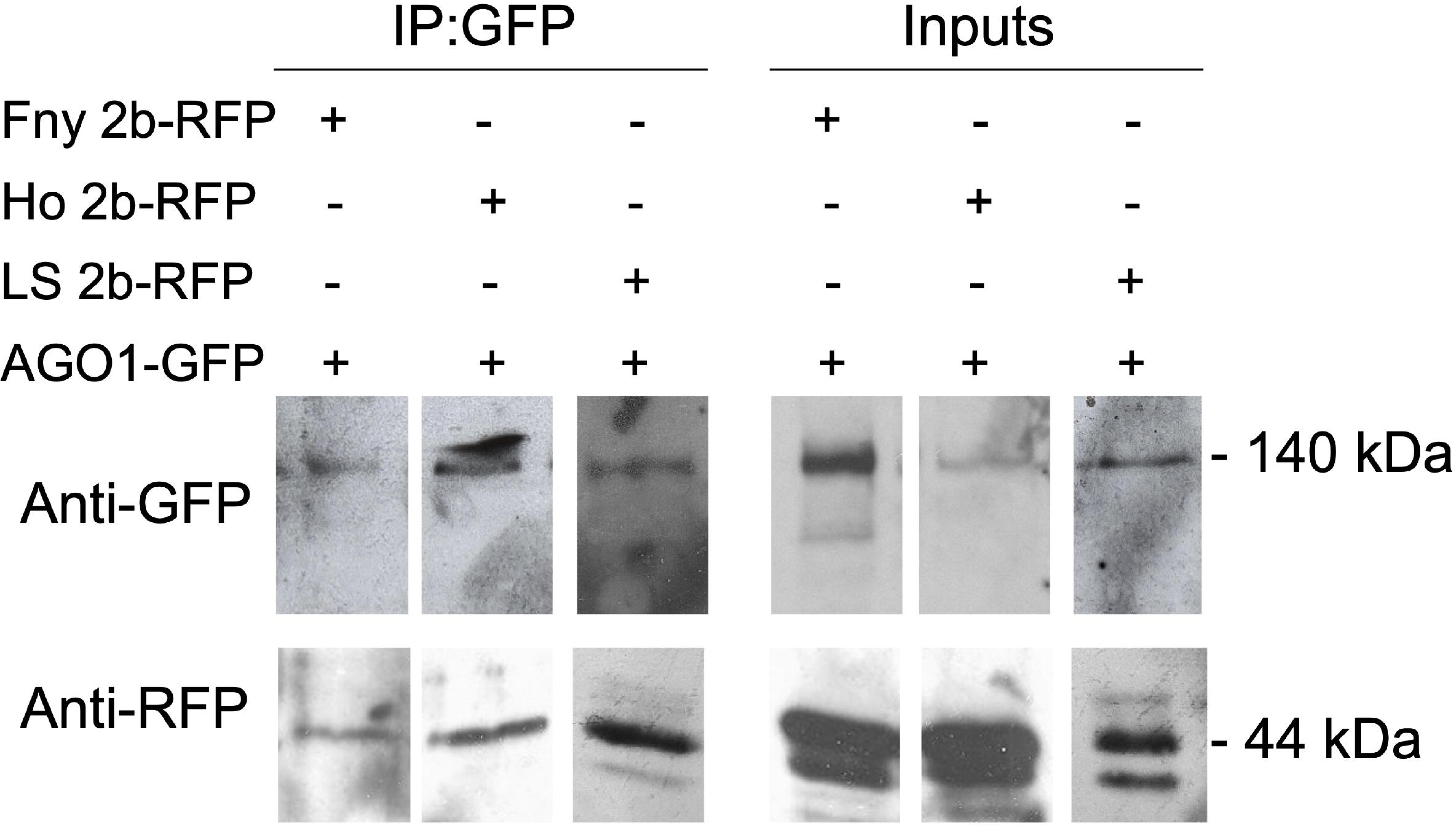
The *Arabidopsis thaliana* Argonaute 1 protein interacts with the 2b proteins of Fny-CMV, Ho-CMV and LS-CMV *in planta*. Proteins were transiently expressed in *N. benthamiana* leaf tissue using agroinfiltration with cells carrying T-DNAs expressing a GFP-AGO1 fusion protein together with cells containing T-DNAs encoding RFP-tagged CMV 2b protein orthologs. For co-immunoprecipitation total protein was extracted from leaf samples and subjected to immunoprecipitation with GFP-Trap beads followed by immunoblot analysis (IP:GFP). For all samples, 2b-RFP was detected in input samples with a corresponding band of approximately 44kDa. Following GFP-pull down, 2b-RFP fusion proteins engineered from the respective 2b orthologs of Fny-CMV (Fny 2b-RFP), Ho-CMV (Ho 2b-RFP) and LS-CMV (LS 2b-RFP) were detected with anti-RFP antibodies. Original blots used to make composite image are shown in Figure S3.

### The Fny-CMV 2b protein inhibits AGO1 RNA endonuclease activity more strongly than either of the 2b proteins encoded by either LS-CMV or Ho-CMV

AGO1 negatively regulates the expression of AGO2 via miR403-directed cleavage of the *AGO2* mRNA (Allen et al., 2005; Lobbes et al., 2006). For example, inhibition of AGO1 activity by the Fny-CMV 2b protein results in an increase in the steady state level of *AGO2* mRNA (Harvey et al., 2011). *AGO2* transcript accumulation was assessed using reverse transcription coupled with a quantitative polymerase chain reaction (RT-qPCR) as a measure of AGO1 inhibition by 2b proteins using primers specific for *NbAGO2* and three housekeeping transcripts (*F-BOX*, *PP2A* and *L23*) (Figure 9; Figure S2).

*NbAGO2* transcript accumulation was consistently increased between four and six-fold in agroinfiltrated leaf patches expressing the Fny-CMV 2b protein. Transient expression of the 2b protein from either LS-CMV or Ho-CMV also both consistently increased *NbAGO2* accumulation, but to a lesser extent (between two and three-fold) than Fny-CMV 2b protein (Figure 9; Figure S2).

**Figure 9.**
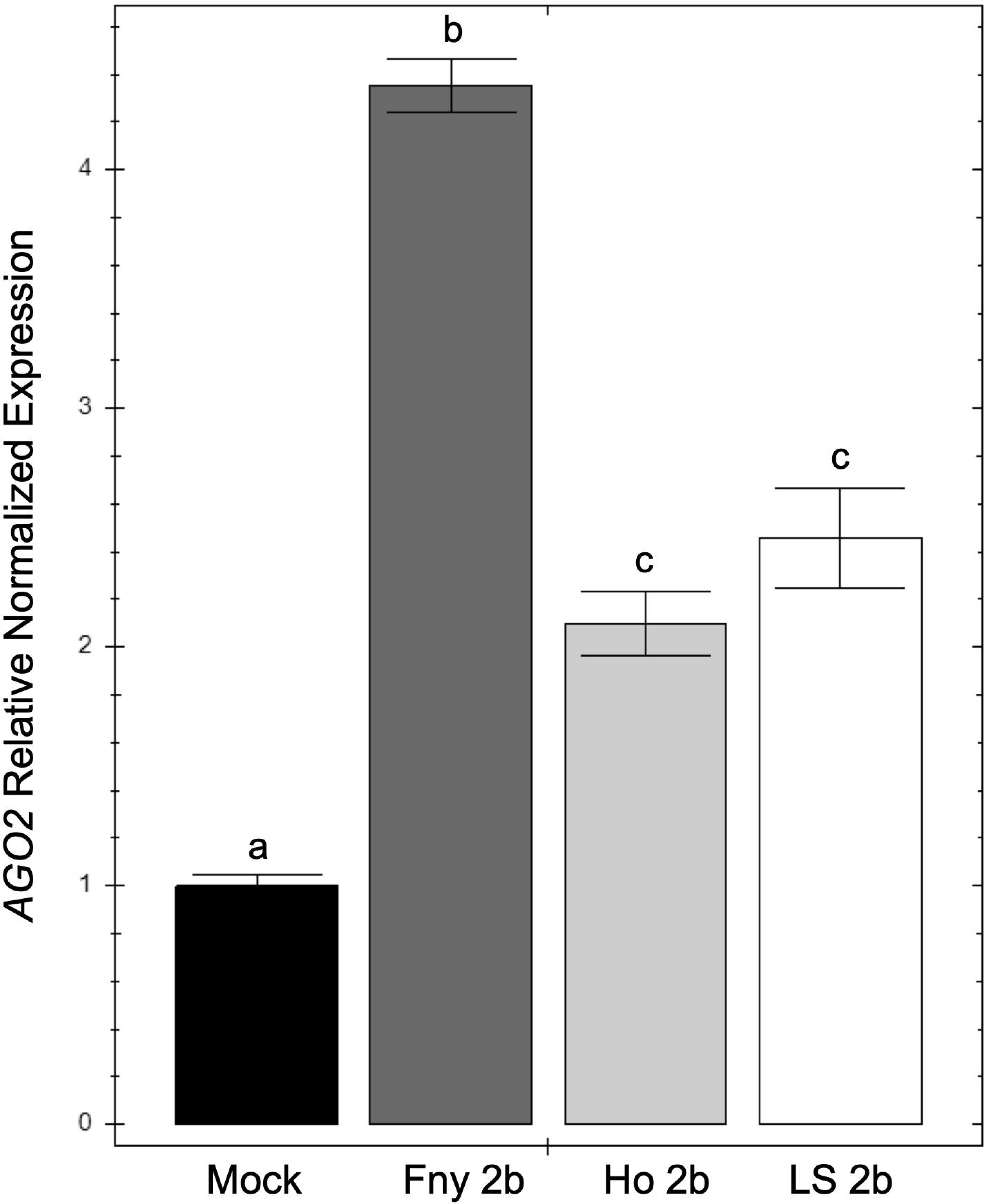
*AGO2* expression increases following transient expression of CMV 2b proteins. Relative levels of *NbAGO2* expression, detected by RT-qPCR in *N. benthamiana* leaves transiently expressing 2b proteins from Fny-CMV, Ho-CMV, or LS-CMV. *AGO2* expression was measured relative to the levels of three *N. benthamiana* housekeeping gene transcripts (*L23*, *PP2A* and *F-BOX*). *AGO2* transcript accumulation was elevated by all CMV 2b protein orthologs, and highest in tissues expressing the Fny-CMV 2b protein. Mock indicates tissue agroinfiltrated with *A. tumefaciens* cells carrying a control ‘empty’ T-DNA. Error bars represent standard error around the mean for three technical replicates. Different lowercase letters indicate statistically significant (α = 0.05) differences in *AGO2* transcript accumulation (analysis of variance and Tukey’s *post hoc* test).

## DISCUSSION

### The relatively lower proportion of Ho-CMV 2b protein that accumulates cytoplasmically may account for its atypical characteristics compared to 2b proteins encoded by other IA Subgroup CMV strains

The localization of the 2b protein is important as, amongst other things, its VSR activity is more active in the cytoplasm (Lewsey et al., 2009; Du et al., 2014a). We found marked differences in the intracellular distributions of three CMV 2b proteins that may explain their different abilities to interact with other factors (such as host AGO1 proteins) or that may be related to symptom induction during CMV infection. The 2b protein from the Subgroup II LS-CMV is mainly limited to the nucleus, while the 2b protein from the Subgroup IA Fny-CMV accumulates in the nucleus and cytoplasm. We found that the distribution between the nucleus and cytoplasm of the 2b protein from Subgroup IA Ho-CMV was intermediate between Subgroup II LS-CMV and Subgroup IA Fny-CMV. Ho-CMV 2b protein could accumulate in both the nucleus and the cytoplasm (like Subgroup IA Fny-CMV) but a much greater proportion partitioned in the nucleus (like Subgroup II LS-CMV).

The phosphorylation status of the 2b protein controls its import and export from the nucleus (Nemes et al., 2017; Kim et al., 2022) and phosphorylation of serine residues 40, 42 and 28 have been shown to influence nuclear localization (Lewsey et al., 2009; Nemes et al., 2017; Kim et al., 2022). Interestingly, certain of the differences between the amino acid sequences of the Ho-CMV 2b and the 2b proteins of other Subgroup IA Fny-CMV and Y-CMV are substitutions of serine with other amino acids (Figure 1). The Fny-2b sequence encodes serine residues at position 47 and 80 and the Y-2b sequence encodes serine residues at position 47 and 77 but in Ho-2b these residues are replaced with alanine, leucine and proline residues (A47, L77 and P80). Additionally, the L77 and P80 substitutions lie within the newly characterized nuclear export sequence between residues (Kim et al., 2022). It has not been confirmed that these serine residues can be phosphorylated, or play roles in 2b protein localization, however, the differences between Ho-CMV 2b protein and orthologs of other Subgroup IA CMV strains suggests a potential explanation for differences in localization and, perhaps, other properties of the Ho-CMV 2b protein.

### Limited interaction of the Ho-CMV and LS-CMV 2b proteins with cytoplasmic AGO1 may explain why they disrupt miRNA-regulated gene expression less than the Fny-CMV 2b protein

The reconstituted Ho-CMV 2b protein interacted directly with AGO1 and interfered with its ability to suppress AGO2 RNA accumulation, contradicting a previous suggestion that this 2b protein is unable to interact with AGO1 (Takahashi et al., 2022). Another surprising finding was that the 2b protein of the Subgroup II strain LS-CMV interacted directly with AGO1 and inhibited AGO1 activity, as indicated by co-localization studies and increased steady-state accumulation of AGO2 mRNA, respectively. Most previous studies on the interactions of 2b proteins of Subgroup II CMV strains such as LS or Q, which have identical amino acid sequences (Mayers et al., 2000), suggest that they have little or no effect on miRNA-regulated changes in gene expression (Chapman et al. 2004; Zhang et al., 2006; Lewsey et al., 2007). This was thought to indicate an inability of LS 2b to inhibit AGO1 due to either a lack of binding site (Lewsey et al., 2007) or reduced stability of the LS-2b protein in vivo (Zhang et al., 2006).

Neither of the 2b proteins from LS- or Ho-CMV inhibit the negative regulation by AGO1 of AGO2 RNA accumulation to the same magnitude as the Fny-CMV 2b protein. Our imaging indicates that although the LS-CMV and Ho-CMV 2b proteins interact with AGO1, they do so predominantly with AGO1 that is present in the nucleus. A possible interpretation of our results is that nuclear-localized AGO1 has a lesser role in miRNA-mediated regulation of RNA accumulation. This fits with the current thinking that degradation of miRNA occurs outside the nucleus (Li et al., 2016; Ma and Zhang, 2018) and supports the proposal that the reduced disruption of miRNA regulation by 2b proteins from Subgroup II strains may be due to their increased nuclear localisation (Du et al., 2014a). Both LS-CMV and Ho-CMV 2b proteins appeared to increase AGO2 levels to some extent which suggests that there may be a threshold of inhibition of AGO1 activity, which is required for a CMV 2b protein to be able to induce a strong change in plant developmental phenotype, as seen in transgenic plants expressing Subgroup IA CMV 2b proteins, or in plants infected by Subgroup IA CMV strains (for example, see Lewsey et al., 2007).

### Interactions between 1a and 2b proteins from different CMV strains may shed light on the mechanism by which CMV modifies plant-aphid relationships

In *A. thaliana*, the interaction of CMV 2b with AGO1 induces feeding deterrence against aphids in infected plants. Our observation that the LS-CMV 2b protein, possibly because of its predominant localization in the nucleus, does not inhibit the AGO1-mediated negative regulation of host transcript accumulation to the same extent as the Fny-CMV 2b protein helps explain why the 2b protein of Fny-CMV but not that of LS-CMV induces strong antibiosis against the aphid *Myzus persicae* (Westwood et al., 2013; Watt et al., 2020). In natural infections of *A. thaliana*, the Fny-CMV 2b protein physically interacts with the Fny-CMV 1a protein, which modulates the 2b-AGO1 protein-protein interaction to prevent the induction of feeding deterrence (Watt et al., 2020). In this study we found that the reconstituted 2b protein of Ho-CMV also interacted with the Fny-CMV 1a protein, but that the LS-CMV 2b did not. It is puzzling why the LS-CMV 1a protein, which shares only 86% amino acid sequence identity with the Fny-CMV or Ho-CMV 1a proteins, is able to interact with the 2b proteins of Ho-CMV and Fny-CMV, while it cannot interact with LS-CMV 2b protein.

Feeding deterrence is also induced in infected tobacco plants but this occurs via a different mechanism whereby the 1a protein induces feeding deterrence and the 2b protein suppresses it. Our observation that the LS-CMV 2b protein does not appear to interact with the CMV 1a proteins of either Fny-CMV or LS-CMV clarifies the mechanism by which the 2b proteins of either LS-CMV or Fny-CMV inhibit the induction of antibiosis against aphids in infected tobacco plants. Infection of tobacco plants with a mutant of Fny-CMV unable to express the 2b protein (Fny-CMVΔ2b) induces antibiosis against *M. persicae* (Ziebell et al., 2011). The viral protein responsible for inducing this anti-aphid resistance in tobacco is the Fny-CMV 1a protein, but induction of resistance can be prevented by either the Fny-CMV 2b protein or by the LS-CMV 2b protein (Tungadi et al., 2020). Arinaitwe and colleagues (2023) proposed alternative hypotheses to explain how 2b proteins impede induction of resistance to aphids in tobacco: through inhibition of jasmonate-dependent signaling (Lewsey et al., 2010; Westwood et al., 2014), or through formation of 2b-1a protein-protein complexes to prevent the 1a protein interacting with a host factor (based on Watt et al., 2020). Since we show here that the LS-CMV 2b protein does not bind to the Fny-CMV 1a protein, it follows that in tobacco plants the 2b proteins of Fny-CMV and LS-CMV must inhibit 1a protein-induced aphid resistance by inhibition of jasmonate-induced defenses.

### Conclusions

Our results suggest that the degree of localization of 2b proteins of different CMV strains in the nucleus versus the cytoplasm is a major determinant of their activity *in planta*. An important consequence of the accumulation of the Fny-CMV 2b protein in the cytoplasm is that its ability to bind to and inhibit AGO1 is stronger than that of the 2b proteins of the Subgroup II strain LS-CMV and of the unusual Subgroup IA strain Ho-CMV, which predominantly accumulate in the nucleus. We found that the modulation of 2b-AGO1 protein-protein interactions by the CMV 1a protein is specific to the 2b proteins of Subgroup IA CMV strains. It appears that the localization of CMV 2b proteins as well as their ability to interact with the 1a protein conditions their ability to induce disease symptoms and modify miRNA-regulated gene expression and, by extension, to modify host-vector interactions.

## MATERIALS AND METHODS

### DNA constructs

Viral sequences used in this study were derived from the Fny strain of CMV (NCBI accession NC002035) (Roossinck and Palukaitis, 1990), the LS strain of CMV (accession AF416900) (Roossinck, 2002) and the Ho strain of CMV (accession LC593245) (Takahashi et al., 2020). A cDNA clone for the 2b protein of Ho-CMV was recapitulated by site-directed mutagenesis of the coding sequence of the Fny-CMV 2b protein using the Q5 Site-Directed Mutagenesis Kit (New England Biolabs). The resultant DNA product encoding the recapitulated Ho-CMV 2b protein contained the amino acid substitutions S47A, V72A, S80P and A106V (Figure 1) and was authenticated by automated Sanger sequencing with both forward and reverse primers (Sanger et al., 1977; Smith et al., 1986) by Source BioScience (Cambridge UK).

Plasmids encoding fluorescently tagged CMV 2b proteins were generated using Gateway^®^ cloning. CMV 2b sequences were amplified with the primers shown in Table S1 to generate the desired *att*B-flanked DNA fragments. The *att*B-flanked fragments were gel purified and cloned into a pDONR221 (Invitrogen) donor plasmid by incubation with BP clonase (Invitrogen) for 1 hour at 25°C. Verified entry clones were used to transfer the CMV 2b sequences into pSITE (Chakrabarty et al., 2007) destination plasmids following incubation with LR clonase (Invitrogen) for 1 hour at 25°C. Sequences encoding Ho-CMV, Fny-CMV and LS-CMV 2b proteins were transferred into pSITE-2NB or pSITE-4NA (Chakrabarty et al., 2007) to yield plasmids expressing Ho 2b-GFP, Fny 2b-GFP and LS 2b-GFP or Ho 2b-RFP, Fny 2b-RFP and LS 2b-RFP, respectively. Additionally, pMDC32-based plasmids were used to express GFP-1a and mRFP-1a and have been previously described (Watt et al., 2020). Sequences encoding Ho-CMV 2b were also transferred into pSITE-cEYFP-N1 or pSITE-nEYFP-N1 (Martin et al., 2009) to generate plasmids expressing YFPc-2bHo or YFPn-2bHo, respectively for BiFC imaging.

Other constructs used for BiFC imaging were based on a pROK background expressing sYFPn-2bFny, sYFPc-2bFny, sYFPn-2bLS, sYFPc-2bLS, sYFPn-AGO1 and sYFPc-AGO1 (González et al., 2010) and sYFPn-1aFny and sYFPc-1aFny (Watt et al., 2020) and have been previously described. The sYFP tags within the pSITE and pROK plasmids were not compatible since the pSITE YFP sequence had been split between residue 174 (Asp) and 175 (Gly) to create N- and C-terminal halves while the pROK plasmids contained a YFP sequence split between residue 155 (Ala) and 156 (Asp) to produce N- and C-terminal halves. To allow BiFC imaging of pSITE and pROK based sYFP tags the pSITE YFPc sequences were extended and pSITE YFPn sequences shortened accordingly using the Q5 Site-Directed Mutagenesis Kit (New England Biolabs) and the primers listed in Table S1.

### Agroinfiltration of Plants

*Nicotiana benthamiana* Domin. seeds were germinated and grown for three weeks in Levington M3 compost (Scotts, Surrey UK) in a Conviron (Manitoba, Canada) growth room maintained at 22°C, 60% relative humidity and 200 μmol.m^-2^.s^-1^ photosynthetically-active radiation under a 16h light - 8h dark cycle. The third or fourth true leaves of the *N*. *benthamiana* plants were used for agroinfiltration assays.

*Agrobacterium tumefaciens* (GV3101) cells harboring plasmids for the expression of fluorescently tagged proteins were cultured overnight at 28°C in 50 ml of liquid LB medium (Sambrook et al., 1989) containing the appropriate plasmid antibiotic and 50 μg/ml rifampicin and 10 μg/ml gentamicin. Cultures were pelleted by centrifugation for 15 min at 5000 *g* and re-suspended in 5 ml MMA buffer (10 mM MgCl_2_, 10 mM MES pH 5.6, and 100 μM acetosyringone). Suspensions were diluted with MMA buffer to an OD_600_ of 0.5 and incubated at room temperature for two hours before being infiltrated into the abaxial side of the leaf using the blunt end of a syringe. Agroinfiltrated *N. benthamiana* plants were returned to the growth room for four days before leaves were removed for imaging, protein extraction or qPCR analysis.

### Fluorescent microscopy

All fluorescent imaging was conducted using a Leica Model SP5 (Leica Microsystems, Heidelberg, Germany) confocal microscope. Fluorescence was monitored from either GFP, mRFP or re-constituted sYFP fluorophores of tagged proteins following excitation at 488 nm, 561 nm or 514 nm, respectively. Nuclear staining was achieved with 300 nM DAPI (Invitrogen) solution infiltrated into the abaxial side of *N. benthamiana* leaves and left for 30 minutes. Excess dye was flushed from the leaf by infiltration with PBS and the leaf was imaged using the UV laser line at 405 nm.

### Western immunoblot analysis

To extract total leaf protein, 100 mg of agroinfiltrated leaf tissue was ground to a powder with a pestle and liquid nitrogen. The powdered sample was homogenized in protein extraction buffer [10% glycerol, 25 mM Tris-HCl (pH 7.5), 200 mM NaCl, 1 mM ethylenediaminotetraacetic acid (EDTA), 0.15% IGEPAL^®^ CA-630 (Octylphenoxy poly(ethyleneoxy)ethanol, branched), 10 mM dithiothreitol, and protease inhibitor cocktail (Roche)] and centrifuged for 4 min at 12,000 *g* at 4°C, to pellet the crude extract and the supernatant collected.

Protein extracts were analysed on an SDS-PAGE gel (Laemmli, 1970) and electrophoretically transferred onto a nitrocellulose membrane (Towbin et al., 1979). For immunological detection of GFP-tagged and RFP-tagged proteins, rabbit anti-GFP (1:1000) and mouse anti-RFP (1:2000) (ChromoTek) monoclonal primary antibodies were used in combination with anti-rabbit or anti-mouse IgG horseradish peroxidase conjugated secondary antibodies. The blotted membrane was incubated with Pierce Enhanced Chemiluminescence ECL-Plus substrate and chemiluminescence signals imaged using X-ray film (Fuji).

### Co-Immunoprecipitation

GFP-tagged or RFP-tagged proteins were immunoprecipitated from total leaf protein extract by incubation with GFP-Trap or RFP-Trap magnetic agarose beads (ChromoTek). The Co-immunoprecipitation assay was carried out according to ChromoTek’s instructions whereby 300 μl of protein extract was diluted with 500 μl of ice-cold dilution buffer [10 mM Tris-HCl (pH 7.5),150 mM NaCl, 0.5 mM EDTA] and incubated with 25 μl of magnetic beads for 1 hr at 4°C. The magnetic beads were washed three times with ice-cold dilution buffer, re-suspended in 50 μl 2 x SDS-sample buffer (Laemmli, 1970) and heated for 10 min at 95°C to dissociate immunocomplexes from the beads. The supernatant was analysed by SDS-PAGE.

### RT-qPCR analysis

Total RNA was extracted from 100mg of frozen leaf tissue using Norgen Total RNA purification kit (Norgen Biotek, ON, Canada), according to the manufacturer’s instruction with the optional addition of 2-mercaptoethanol (0.01%). RNA concentrations and sample purity were measured using a Nanodrop ND-2000 spectrophotometer (Nanodrop Technologies, Rockland, DE, USA) and only samples with an A260/A280 ratio between 1.8-2.0 were used for subsequent analysis. First strand cDNA was synthesised from 500 ng of total RNA using the GoScript^TM^ Reverse Transcription System and oligo (dT)_15_ primers (Promega Madison, WI, USA) according to their instructions with the optional addition of RiboLock RNase inhibitor (20 units) (Thermo Fisher Scientific, MA, USA).

RT-qPCR was performed in 96-well plates using the CFX96 real-time PCR detection system (Bio-Rad, Hercules, CA, USA). Six biological replicates were used from each treatment and each qPCR run used three technical replicates of each sample (i.e., cDNA from the same RNA preparation). Each 10 μl reaction consisted of 5 μl 2X SYBR green, 1 μl cDNA, 1.25 μl forward and reverse primers (3 pmol/ml) and 1.5 μl DNAse free water. The thermal profile of the reaction consisted of an initial 10-minute denaturation at 95°C, followed by 40 cycles of 10 seconds at 95°C and 10 seconds at 60°C with fluorescence acquisition after each cycle. Finally, a dissociation curve was generated by increasing temperature from 65°C to 95°C, to verify primer specificity.

*PP2A*, *L23* and *F-BOX* were used in combination as reference genes since they have stable constitutive expression in *N. benthamiana* that is unaffected by virus infection (Liu et al., 2012). *AGO2* expression was then normalised relative to the mean of *PP2A*, *L23* and *F-BOX* expression for each sample. Relative quantification was performed using the Bio-Rad CFX Manager software (version 1.1) with default parameters. Primer sequences used for *PP2A*, *L23*, *F-BOX* and *AGO2* amplification followed those used in a previous study in *N. benthamiana* (Liu et al., 2012) and are shown in Table S1.

## Acknowledgements

We thank Adrienne E. Pate for expert technical assistance, Tomás Canto for provision of DNA constructs and Satish Bharathwaj Viswanathan for useful discussions. Work was funded by a grant from the UK Biotechnological and Biological Sciences Research Council (BB/P023223/1). SC and LGW received studentships from the BBSRC-Cambridge University Doctoral Training Program (BB/M011194/1). AMM was supported by a BBSRC grant (21ROMITIGATIONFUND CAMBRIDGE BB/W510609/1) and a grant from The Leverhulme Trust (RPG-2022-134).

## Supplementary data

**Figure S1.**
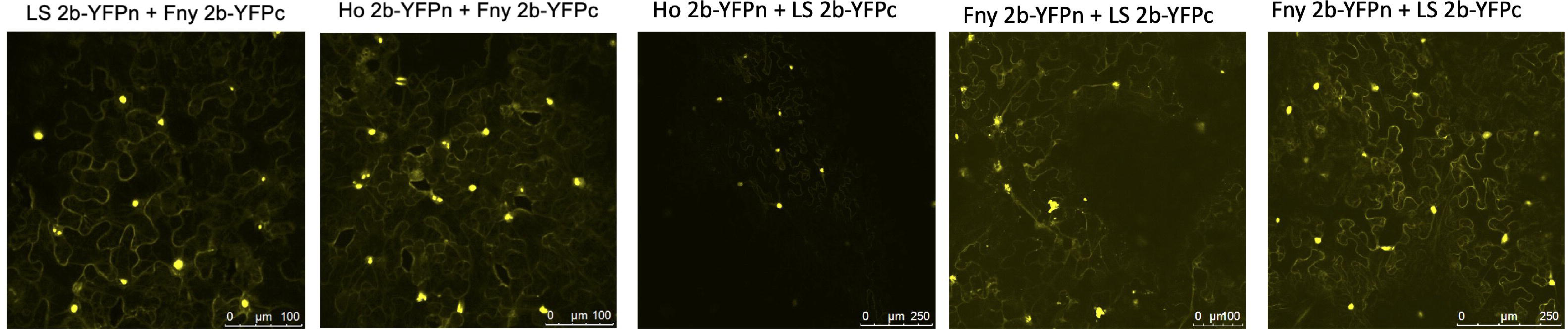
Subcellular localization of cucumber mosaic virus 2b proteins from different CMV strains using reciprocal combinations of BiFC constructs. Bimolecular fluorescence complementation (BiFC) was used to compare the self-interaction properties of the 2b proteins of Fny-CMV, LS-CMV, and Ho-CMV, using fusion proteins with the N- and C-terminal domains of the yellow fluorescent protein (2b-YFPn and 2b-YFPc). The 2b proteins of all three strains formed heterodimers in vivo, indicated by yellow fluorescence. The intracellular distributions of these heterodimers were consistent with those seen in Figure 2C for reciprocal combinations of 2b-YFPn or 2b-YFPc fusion proteins.

**Figure S2.**
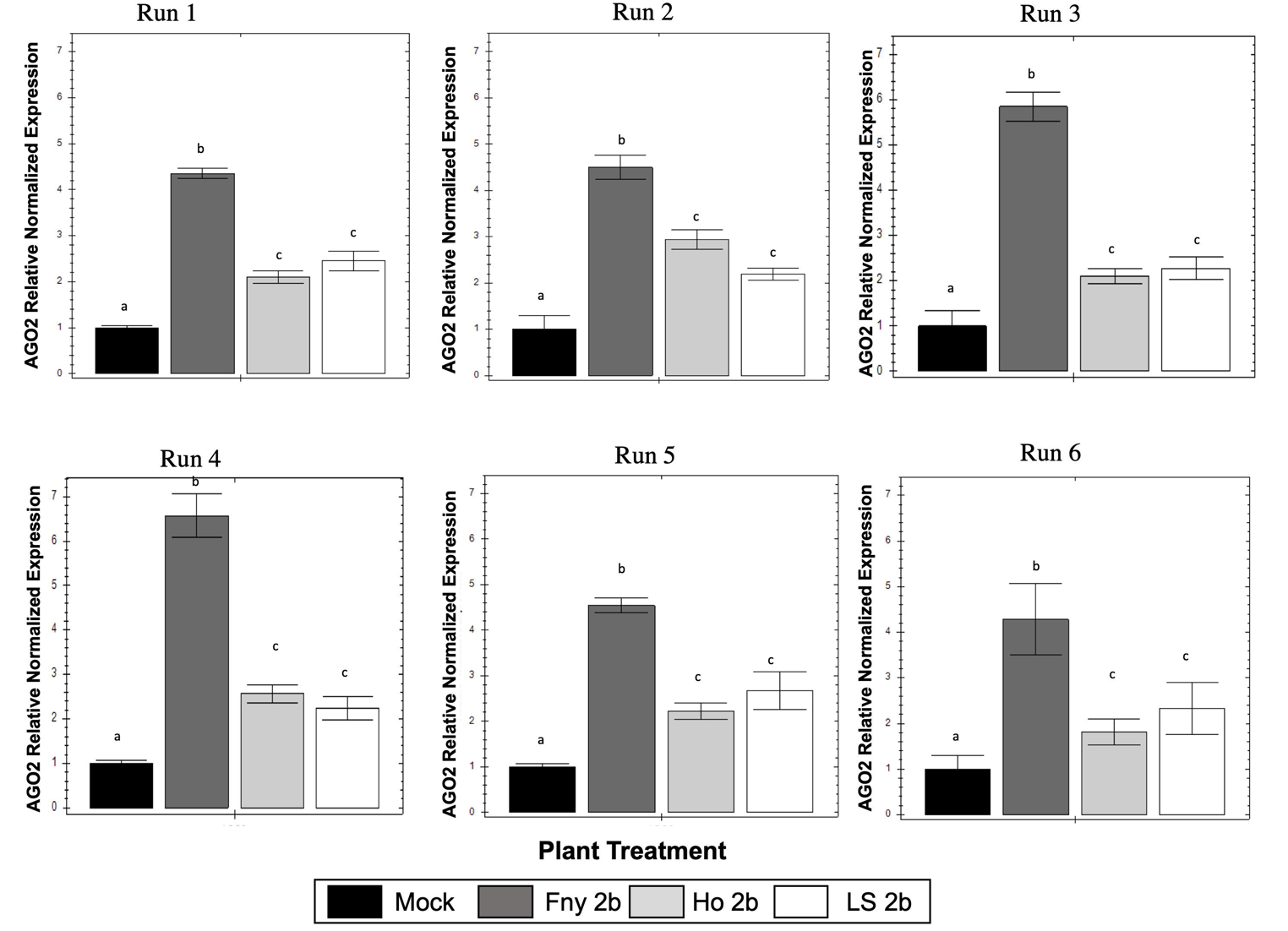
Analysis of *NbAGO2* expression. Altogether six independent agroinfiltration experiments were carried out to examine the inhibitory effects of the 2b proteins of Fny-CMV, Ho-CMV, or LS-CMV on AGO1. Accumulation of *AGO2* mRNA was chosen as the proxy for this since its turnover is mediated by miR403-directed AGO1 endonuclease activity. An increase in the steady-state level of *AGO2*, measured using RT-qPCR, indicates that inhibition of AGO1 activity has occurred. Error bars represent standard error around the mean for three technical replicates. Different lowercase letters indicate statistically significant (α = 0.05) differences in *AGO2* transcript accumulation (analysis of variance and Tukey’s *post hoc* test). Mock: *AGO2* accumulation in tissues infiltrated with *A. tumefaciens* cells transformed with a control (‘empty’) T-DNA vector.

**Figure S3.**
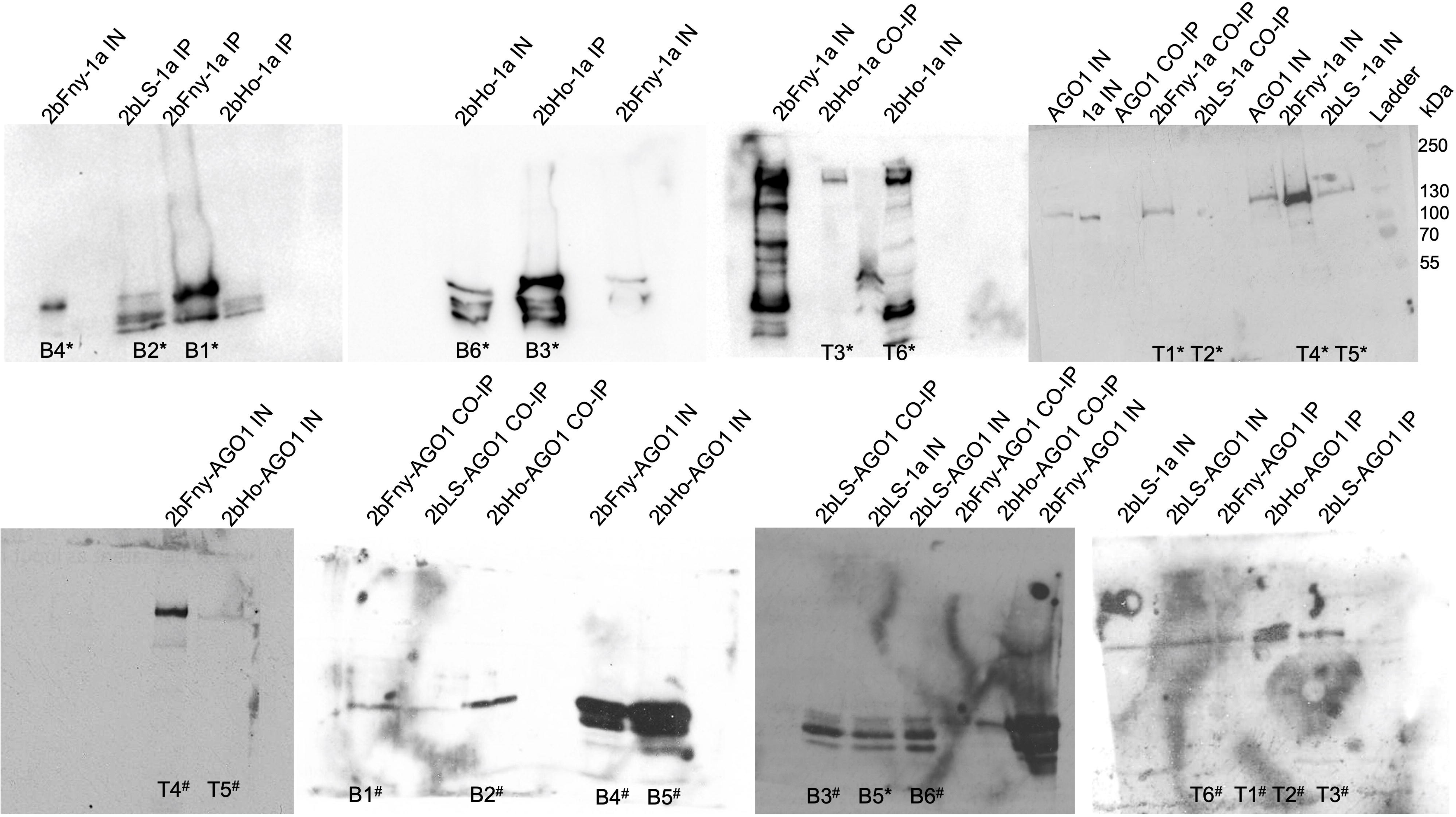
Whole gel images showing association of 2b proteins from the Fny, Ho and LS strains of CMV with AGO1 or 1a proteins demonstrated by co-immunoprecipitation. RFP-tagged 2b proteins from Fny-CMV, Ho-CMV or LS-CMV were co-expressed with GFP tagged AGO1 or 1a proteins in *N*. *benthamiana* leaves. Total protein was subjected to immunoprecipitation with GFP-Trap or RFP-Trap beads followed by immunoblot analysis with anti-RFP antibodies to detect 2b-RFP or anti-GFP antibodies to detect GFP-1a or GFP-AGO1. After Immunoprecipitation of 2b-RFP with RFP-trap beads, GFP-1a only be detected when co-expressed with 2b-RFP from Fny-CMV and Ho-CMV but not LS-CMV. After Immunoprecipitation of AGO1-GFP with GFP-trap beads, 2b-RFP was detected from Fny-CMV, Ho-CMV and LS-CMV. Bands are labelled with the contents of the total protein extract and treatment: input sample (IN), immunoprecipitation with Trap antibodies (IP) or Co-immunoprecipitation with interacting partner (CO-IP). The position where each band appears in the composite blot figures is denoted by T or B referring to the top or bottom row of gel strips. Asterisks (*) relate to bands used to form the composite blot in Figure 5 and hashtags (#) relate to bands used to form the composite blot in Figure 8.

