## Supplemental Table S1 for "Strain-specific differences in the interactions of the cucumber mosaic virus 2b protein with the viral 1a and host Argonaute 1 proteins"

**Table S1.** Primer sequences.

| | |
|---|---|
| **Primers for detection of transcripts** | |
| Primer set/transcript name | 5’-3’ Sequence Forward/Reverse |
| *F-BOX* | GGCACTCACAAACGTCTATTTC / ACCTGGGAGGCATCCTGCTTAT |
| *L23* | AAGGATGCCGTGAAGAAGATGT / GCATCGTAGTCAGGAGTCAACC |
| *PP2A* | GACCCTGATGTTGATGTTCGCT / GAGGGATTTGAAGAGAGATTTC |
| *NbAGO2* | TTTCCGTGATGGTGTGAGT / GAGTGATTGCTGGTCGATA |
| **Primers for Q5 Mutagenesis of the Fny-CMV *2b* gene to recapitulate the Ho-CMV *2b* gene** | |
| Primer set name | 5’-3’ Sequence Forward/Reverse |
| 2bA106V | AACGAATGGGTGGAAGGTGCT / ACCGGCGAACCAATCTGTAT |
| 2bS80P | GTTACCCGAGCCTGAGGCCTC / TCCGCCACGTTCACATGG |
| 2bV72A | TGCCGCCATGCGAACGTGGCG / TGACCCTGTCAGTTCCGAACC |
| 2bS47A | GAGAGCGCGTGCAAATCTCAG / TCGCTGGGACTTTTGTGA |
| **Primers for Q5 mutagenesis of *YFP* sequences** | |
| Primer set name and use | 5’-3’ Sequence Forward/Reverse |
| PSITE-YFPC extension | CTTCAAGATCCGCCACAACATCGAGGACGGCAGCGTGCAGCTCGCC / TTCACCTTGATGCCGTTCTTCTGCTTGTCCCCTATCACCACTTTGTACAAGAAAGCTGAACG |
| PSITE-YFPN deletion | GGCAGCGTGCAGCTC / GGCCATGATATAGACGTTGTGG |
| **Primers for Gateway Cloning** | |
| Primer set name | 5’-3’ Sequence Forward/Reverse |
| 2b attB | GGGGACAAGTTTGTACAAAAAAGCAGGCTTCATGGAATTGAACGTAGGT / GGGGACCACTTTGTACAAGAAAGCTGGGTTTCAGAAAGCACCTTCCGC |
